# The periplasmic substrate-binding protein ThuE is required for sucrose uptake and multiplication of *Pseudomonas syringae* pv. tomato DC3000 in tomato leaves

**DOI:** 10.64898/2026.10.25.761734

**Authors:** Chen Deng, Ji-Yun Kim

## Abstract

Phytopathogenic bacteria depend on host-derived carbon for successful colonization of plant tissue. While sucrose is among the most abundant sugars in the leaf apoplast, the steps that limit its acquisition by *Pseudomonas syringae* are not known. In this study, we used base editing to generate mutants of six candidate sucrose utilization genes in *P. syringae* pv. tomato DC3000 (*Pto*). Loss of either the sucrose hydrolase ScrB or both levansucrases had no detectable effect, whereas the growth defect in the loss of the outer membrane porin mutant *scry* was overcome by raising external sucrose. Loss of the periplasmic substrate-binding protein ThuE impaired growth both *in vitro* and *in planta*, reducing bacterial populations to approximately two orders of magnitude lower in tomato leaves. Sucrose import through a single non-redundant inner membrane ABC transporter is therefore required for *Pto* multiplication *in planta* and serves as a potential target for controlling bacterial speck disease.

**Author Summary:** Base editing of six candidate genes shows that *Pseudomonas syringae* pv. tomato DC3000 imports sucrose mainly via an ABC transporter, a step required for bacterial growth in tomato leaves.

## Introduction

Phytopathogenic bacteria have evolved sophisticated molecular mechanisms to recognize and utilize host-derived nutrients, a prerequisite for establishing a nutrient-rich niche (Anderson, 2023; Roussin-Léveillée et al., 2024). Successful colonization of plant tissues inherently relies on the bacterial capability to sense specific chemical cues within the complex host microenvironment through specialized chemoreceptors and two-component regulatory systems. For example, fructose recognition triggers the downstream virulence pathways, including the type III secretion system (Anderson, 2023). By manipulating host physiology, phytopathogenic bacteria successfully convert the intercellular spaces, primarily the apoplasm, into a nutrient-rich environment that supports rapid multiplication.

The exploitation of host-derived sucrose contributes to fitness, growth, and virulence across several distinct *Pseudomonas syringae* (*P. syringae*) pathovars. Recent pan-genome analyses indicate a high conservation of the sucrose utilization (*scr*) gene cluster across diverse geographical strains and pathovars (Dhaouadi et al., 2025). In *P*. *syringae* pv. aesculi (*Psa*), the outer membrane sucrose porin ScrY mediates extracellular sucrose uptake and supports multiplication in woody tissues and foliar environments and the intracellular sucrose hydrolase (invertase) ScrB is required for sucrose catabolism in woody tissues (Dhaouadi et al., 2025). In *P*. *syringae* pv. *syringae* B728a, the disruption of the *scrB* moderately reduces bacterial fitness in the leaf apoplasm (Helmann et al., 2019). Similarly, *Pto* catabolizes sucrose present within the tomato leaf apoplastic fluid (McCraw et al., 2016).

Beyond serving as a primary carbon and energy source, sucrose acts as a signaling molecule in the apoplasm during the infection process (Huynh et al., 1989). In hypersensitive response and pathogenicity (*hrp*) gene-derepressing minimal media designed to mimic the osmotic and nutritional conditions of the apoplasm, the extracellular sucrose induces transcription of bacterial virulence genes, including *hrp* regulators, in *Pto* (Stauber et al., 2012; Turner et al., 2020). Concurrently, the host plant reacts dynamically to bacterial invasion; for instance, tomato sucrose transporter 2 (*Sl*SUT2) promotes *Pto* proliferation through a dual mechanism, inducing sugar efflux into the apoplast while repressing the biosynthesis of salicylic acid, a key plant defence hormone responsible for systemic acquired resistance (Chen et al., 2026).

While the importance of sucrose is increasingly clear, the genetic architecture supporting the uptake and cleavage of sucrose in *P. syringae* has not been resolved. Therefore, the primary objective of this study is to identify *Pto* genes required for sucrose transport and metabolism, and to test whether sucrose acquisition limits bacterial multiplication in tomato leaves.

## Results

### Identification and base-editing mutagenesis of genomic components governing sucrose utilization in *Pto*

To identify genetic determinants governing sucrose utilization in *Pto*, we analyzed the *Pto* genome and identified a putative *scr* gene cluster between nucleotides 960,352 and 968,939 of the chromosome. The *scr* gene cluster comprises a LacI-family transcriptional regulator, a sucrose hydrolase (*scrB*), four trehalose/maltose ABC transport system components (*thuE*, *thuF*, *thuG*, and *thuK*), and a sucrose porin (*scrY*) (Fig. 1A). Apart from the putative *scr* gene cluster, we identified loci encoding levansucrases (*lsc1* and *lsc2*) and phosphotransferase system (PTS) components (Fig. 1A). To test the roles of the PTS component protein-N(pi)-phosphohistidine--D-fructose phosphotransferase (EIIBC), Lsc1, Lsc2, ScrB, ScrY, and ThuE in sucrose utilization, we introduced premature stop codons into their open reading frames by base-editing (Fig. 1B) (Chen et al., 2018).

**Fig. 1.**
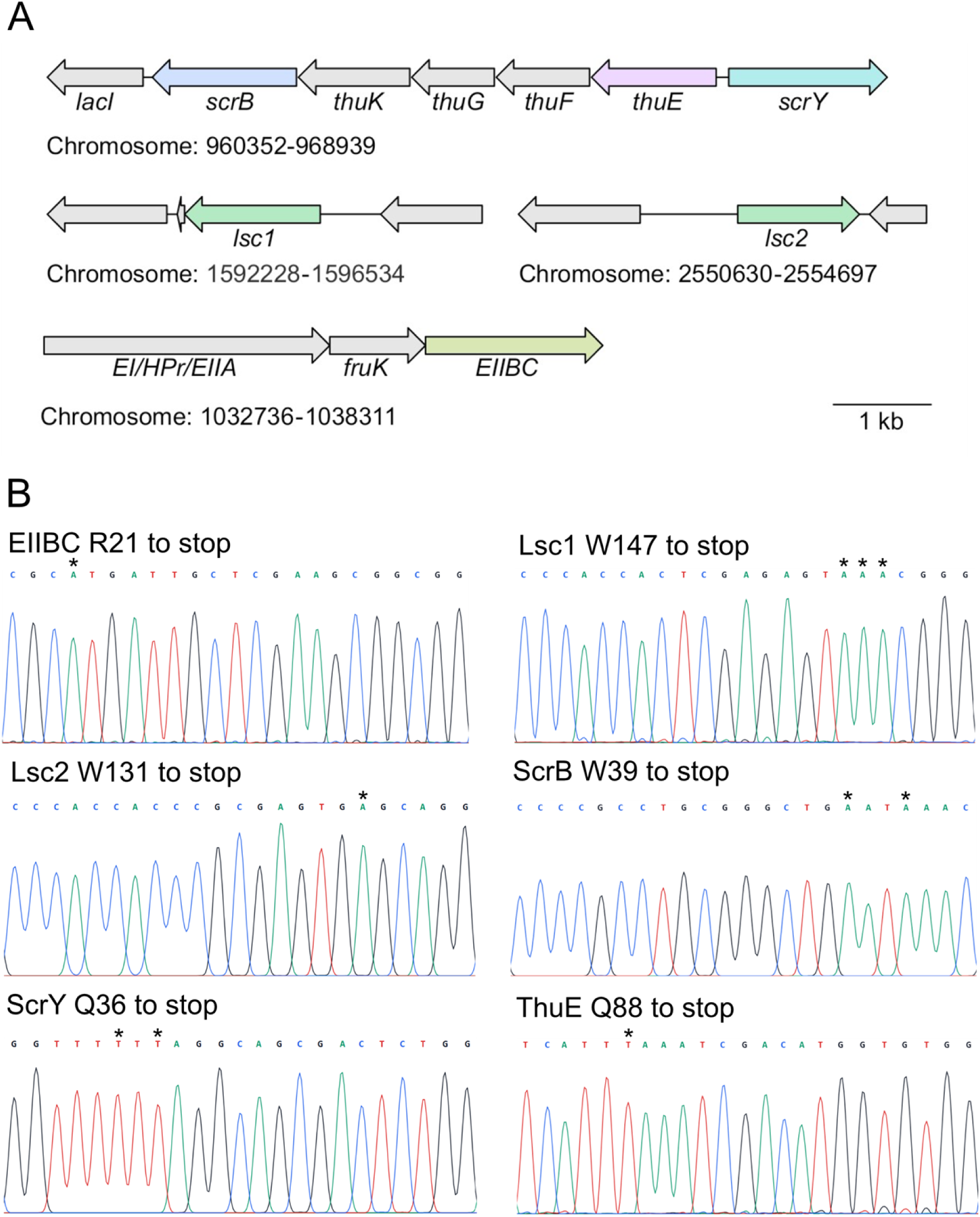
Genomic organization of the putative sucrose utilization genes and verification of the base-edited mutants. (A) Schematic representation of the sucrose utilization genes. *lacI*, LacI family transcriptional regulator. *scrB*, sucrose hydrolase. *thuK*, trehalose/maltose ABC transporter ATP-binding protein. *thuG* and *thuF*, trehalose/maltose ABC transporter permease. *thuE*, trehalose/maltose ABC transporter periplasmic substrate-binding protein. *scrY*, sucrose porin. *lsc1*, levansucrase 1. *lsc2*, levansucrase 2. *EI/HPr/EIIA*, phosphoenolpyruvate-protein phosphotransferase. *fruK*, 1-phosphofructokinase. *EIIBC*, protein-N(pi)-phosphohistidine--D-fructose phosphotransferase. (B) Sanger sequencing chromatograms of EIIBC R21 to stop, Lsc1 W147 to stop, Lsc2 W131 to stop, ScrB W39 to stop, ScrY Q36 to stop, and ThuE Q88 to stop are shown. Base edited positions are marked by asterisk.

In the minimal M9 medium supplemented with limiting sucrose concentration (1 mM), both the sucrose porin mutant *scry* and the periplasmic substrate-binding protein mutant *thue* reached lower optical densities compared to the parental wild-type (WT) strain (Fig. 2A). At the higher sucrose concentration (10 mM), the growth defect of *scry* was markedly minimized (Fig. 2B). This concentration-dependent recovery implies that high external availability of sucrose can bypass the ScrY-dependent outer membrane step, potentially via passive diffusion or through alternative, low-affinity transport pathways such as TonB-dependent receptors (TBDRs). In contrast, *thue* maintained a severe growth defect even at 10 mM sucrose, indicating that the ThuE-dependent ABC transport system is required for efficient sucrose transport across the inner membrane (Fig. 2B).

**Fig. 2.**
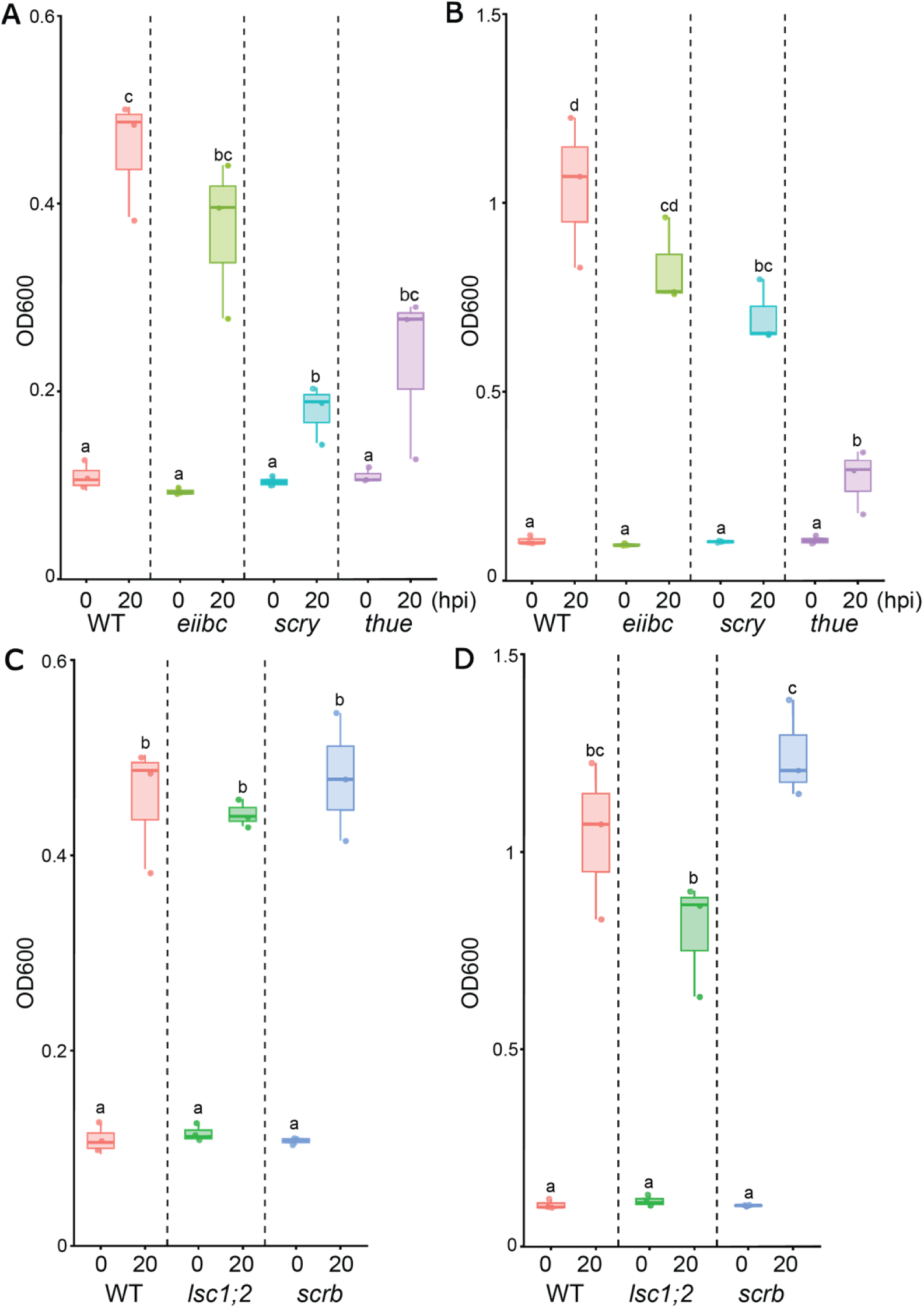
Disruption of *thuE* and *scrY* decreases *Pto* growth in M9 medium supplemented with sucrose as the sole carbon source. (A, B) Growth of the membrane transport defective mutants *eiibc*, *scry*, *thue*, and WT in M9 medium supplemented with 1 mM (A) or 10 mM (B) sucrose. (C, D) Growth of the sucrose metabolism defective mutants *lsc1;2*, *scrB*, and WT in M9 medium supplemented with 1 mM (C) or 10 mM (D) sucrose. Boxplots show the minimum (lower end of the whisker), first quartile (lower edge), median (central line), third quartile (upper edge), and maximum (upper end of the whisker) optical densities at 600 nm (OD600) at 0 and 20 h post-inoculation (hpi). Data are derived from three independent trials, with 0 and 20 hpi measurements paired from the same culture within each trial. Statistical significance was determined using Two-way Repeated Measures ANOVA followed by Tukey’s multiple comparisons test. Groups with different lowercase letters (a, b, c, d) indicate statistically significant differences (*p* < 0.05).

The protein-N(pi)-phosphohistidine--D-fructose phosphotransferas mutant *eiibc* retained a biomass gain comparable to the WT control at both sucrose concentrations (Fig. 2A-B). Because EIIBC is clustered with a fructokinase gene and is likely a fructose-specific PTS component, the absence of a growth defect on minimal M9 medium supplemented with 1 or 10 mM of sucrose argues against the PTS component acting as an alternative route for inner membrane sucrose transport in *Pto* (Fig. 1A).

Together, our *in vitro* growth assay demonstrate that inner membrane sucrose transport in *Pto* relies mainly on the ThuE-dependent ABC transport system.

### Parallel metabolic pathways governed by ScrB and Lsc for *Pto* sucrose metabolism

To evaluate the metabolic processing of sucrose by *Pto*, we characterized *Pto* strains deficient in key sucrose-cleaving enzymes: the sucrose hydrolase ScrB (*scrb*) and the levansucrases Lsc1 and Lsc2 (*lsc1;2*). Protein sequence alignment of *Pto* ScrB with characterized bacterial sucrose-6-phosphate hydrolases from *Lactobacillus gasseri* 224-1 (S6PH) and *Priestia megaterium* (SacAP1) and with sucrose hydrolases from *Priestia megaterium* (SacAP2 and SacAP3) indicates that *Pto* ScrB carries the residues that distinguish sucrose hydrolases from sucrose-6-phosphate hydrolases (Fig. S1) (Dobrange et al., 2024). Both mutants *scrb* and *lsc1;2* maintained biomass gains that were statistically indistinguishable from the WT strain at either 1 or 10 mM of sucrose (Fig. 2C-D). This phenotypic resilience is consistent with *Pto* may flexibly deploy either Lsc or ScrB to cleave sucrose into glucose and fructose.

### Impaired sucrose uptake compromises *Pto* in planta multiplication

Because *thue* exhibited defects in utilizing sucrose *in vitro*, we asked whether sucrose uptake limits bacterial multiplication within the natural host tissue. We determined the viable populations of WT and *thue* in *Solanum lycopersicum* (tomato) M82 leaves via CFU enumeration. When inoculated individually into tomato leaves, the viable population size of *thue* was significantly reduced by 3 dpi. Specifically, the population of *thue* was approximately two orders of magnitude lower (1/100th) than those achieved by the virulent WT *Pto* strain and inoculated leaves showed reduced speck symptoms (Fig. 3). The severe colonization defect of *thue* indicates that the ThuE-dependent sucrose transport system is required for *Pto* to multiply in the tomato leaf apoplast.

**Fig. 3.**
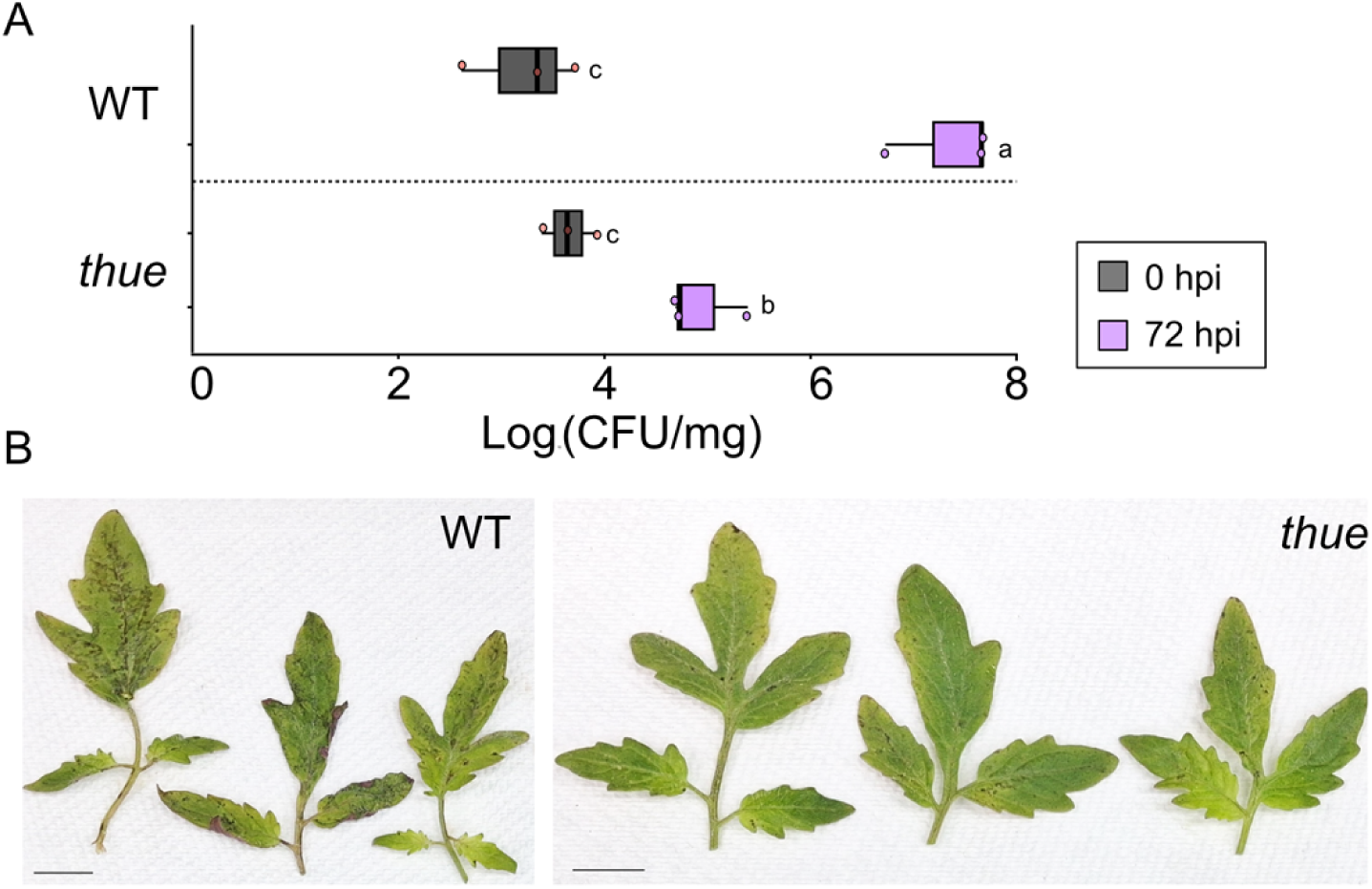
The periplasmic substrate-binding protein ThuE is required for *Pto* proliferation in tomato M82 leaves. **(A)** Tomato M82 plants were dip-inoculated with 2.5 x 10^7^ colony-forming units (CFU) mL^-1^ of wild-type strain (WT) and the *thue* mutant. Boxplots show the minimum (lower end of the whisker), first quartile (lower edge), median (central line), third quartile (upper edge), and maximum (upper end of the whisker) Log_10_ (CFU/mg fresh weight). For samples collected right after inoculation (0 hpi), each data point corresponds to a biological replicate, with three replicates per bacterial strain, where each replicate consists of cotyledons pooled from three individual plants. For samples collected at 72 hpi, each data point represents a biological replicate, with three replicates analyzed per bacterial strain. Each replicate consisted of true leaves sourced from three individual plants. Values with different lowercase letters (a, b, c) are statistically significantly different (*p* < 0.05) by Two-way ANOVA. (B) Representative images of the infected M82 tomato leaves at 72 hpi. Scale bars: 1 cm. This experiment was conducted three times with comparable results.

## Discussion

Among economically important phytopathogenic bacteria, *Pseudomonas syringae* species complex encompasses a widespread and devastating group that infects a wide range of crops while serving as a well-established model for studying plant-microbe interactions (Mansfield et al., 2012). Depriving these pathogens of host-derived carbon is an attractive control strategy, and our results identify the step in the sucrose utilization pathway where such deprivation could not be bypassed.

The recovery of *scry* growth at 10 mM sucrose concentrations indicates that alternative outer membrane translocation pathways, or extracellular sucrose catabolism, operate when sucrose is abundant (Fig. 2A-B). A TBDR mediated route is possible, mirroring the strategy observed in *Xanthomonas campestris* pv. *campestris* (*Xcc*) where the TBDR SuxA-mediated transport across the outer membrane is redundant for extracellular sucrose utilization but remains indispensable for full pathogenesis (Blanvillain et al., 2007). As similar TBDRs may be encoded within the *Pto* genome, redundancy at the outer membrane may allow *Pto* to maximize carbon influx when transient, nutrient-rich microenvironments are encountered within the host tissue.

Phytobacterial pathogens use diverse inner membrane systems to facilitate sucrose translocation. The major facilitator superfamily H^+^-symporter SuxC is required for both sugar utilization and pathogenesis in *Xanthomonas oryzae* pv. *oryzae* (*Xoo*) PXO99 (Zöllner et al., 2026), whereas *Ralstonia solanacearum* UW551 uses ScrA (EIIBC) to allow inner-membrane sucrose transport (Jacobs et al., 2012). In *Pto*, however, the growth defect of *thue* at both 1 and 10 mM sucrose indicate that the ThuE-dependent ABC transport system is the rate-limiting step across the concentrations tested (Fig. 2A-B). Such a transporter may be favoured where apoplastic sucrose is dilute, since binding-protein-dependent systems operate at high affinity, although why *Pto* uses this route rather than an MFS symporter or a PTS remains to be established.

Because ABC transporters import intact sucrose, a dedicated sucrose hydrolase, rather than a sucrose-6-phosphate hydrolase, is expected, and the classification of ScrB as an enzyme targeting unmodified sucrose is consistent with ATP-driven ABC transport, although the assignment remains a prediction from sequence as ScrB activity was not assayed directly (Fig. S1). While disruption of sucrose hydrolases in *Psa*, *Xcc*, and *Xoo* impairs extracellular sucrose utilization, *scrb* grew as well as WT, as did the *lsc1;2* (Fig. 2C-D) (Blanvillain et al., 2007; Dhaouadi et al., 2025; Zöllner et al., 2026). Here, the simplest interpretation is that either enzyme can compensate for the loss of the other. Yet, as the two enzymes act in different compartments (Lsc extracellularly and ScrB in the cytoplasm), such compensation would require extracellular cleavage to feed the cell through hexose uptake, and a *scrb lsc1;2* triple mutant would be required to exclude a further, unidentified sucrose metabolism pathway.

The apoplast of host leaves is a restrictive niche in which successful colonization depends on a pathogen’s capacity to acquire nutrients rapidly while evading or suppressing early host immunity. Against that background, the consequence of losing ThuE *in planta* is substantial: *thue* populations were almost three orders of magnitude below WT at 3 dpi, with correspondingly reduced speck symptoms (Fig. 3A, B). A transport bottleneck defined in minimal medium therefore translates into a colonization defect in the host, which is the clearest evidence that *Pto* depends on apoplastic sucrose. Two limitations bound this conclusion. First, *thue* was neither complemented nor recovered as an independent allele, and cytidine base editors can introduce off-target transitions. Second, growth was tested only on sucrose, so a sucrose-specific uptake defect cannot yet be distinguished from a broader one; this matters because ThuE orthologues are annotated as trehalose/maltose-binding proteins, and in *Sinorhizobium meliloti* a *thuE* mutant grows normally on sucrose (Jensen *et al.*, 2002).

In summary, our study provides a framework for understanding sucrose utilization in *Pto*. Entry through the outer membrane porins and sucrose cleavage tolerate the loss of a single component whereas the ThuE-dependent inner membrane sucrose transport is a non-redundant bottleneck, whose disruption substantially reduces bacterial multiplication in tomato leaves. These findings identify the inner membrane ABC transport machinery as a candidate target for developing anti-virulence strategies against bacterial speck disease.

## Materials and Methods

### Generation of *Pto* nonsense mutants by base editing

To introduce premature stop codons into the coding regions of *scrB* (PSPTO_0885), *scrY* (PSPTO_0890), *lsc1* (PSPTO_1453), *lsc2* (PSPTO_2305), *thuE* (PSPTO_0889), or *EIIBC*(PSPTO_0956), the CRISPR (clustered regularly interspaced short palindromic repeats) Guide ribonucleic acid (RNA) Design tool available on Benchling.com was utilized to select guide RNA target sites.

Two synthesized 24 nucleotide complementary DNA oligonucleotides (Bionics) were annealed to form a double-stranded DNA oligonucleotide (dsDNA) after T4 polynucleotide kinase-mediated phosphorylation for creating CRISPR RNA (crRNA) protospacer sequences. Annealed dsDNA carried a 5′ overhang of GTGG on the sense strand and a 5′ AAAC overhang on the antisense strand (Table S1). To construct the bacterial base-editing vector, annealed dsDNA was separately incorporated into the vector pnCasPA-BEC utilizing the Golden Gate Assembly method (Chen et al., 2018).

To prepare *Pto* competent cells, *Pto* glycerol stock was used to inoculate 1.5 mL of 2x yeast extract tryptone (dYT) medium [tryptone at 16 g L^-1^ , yeast extract at 10 g L^-1^ , NaCl at 5 g L^-1^ , pH = 7] containing 50 μg mL^-1^ rifampicin. The overnight *Pto* culture was harvested by centrifugation (8000 g, 2 min, 22 °C), washed in 1 mL of 0.3 M sucrose (8000 g, 2 min, 22 °C), and the resulting bacterial pellet was resuspended in 50 μL of 0.3 M sucrose. 500 ng of pnCasPA-BEC-sgRNA was introduced into *Pto* by electroporation using a 2 mm cuvette with parameters set at 2000 voltage. Following electroporation, *Pto* was allowed to recover in 500 μL of dYT medium for 2 hours at 28 °C with shaking at 200 revolutions per minute (rpm) before plating on dYT agar containing 50 μg mL^-1^ rifampicin and 10 μg mL^-1^ gentamicin for selection. To verify mutations in the target genes, gene-specific primers were employed to amplify the regions of interest (refer to Table S1), and the resulting amplicons were subjected to analysis via Sanger sequencing (Bionics). pnCasPA-BEC-sgRNA was cured from *Pto* by selecting viable cells grown on King’s medium B (KB) agar containing 50 μg mL^-1^ rifampicin and 5 % sucrose.

### *In vitro* growth assay of *Pto*

Glycerol stocks maintained at -80°C were used to inoculate KB agar plates supplemented with 50 μg mL^-1^ rifampicin. All *Pto* strains were incubated at 22°C for a duration of two days. For the preparation of starting cultures, the *Pto* and its mutant strains grown on KB agar were resuspended in a 10 mM MgCl_2_ solution. The bacterial cell density was adjusted to an optical density 600 (OD600) of approximately 0.1 in 500 μL of modified M9 minimal medium [42.2 mM Na_2_HPO_4_, 22 mM KH_2_PO_4_, 23.5 mM (NH_4_)_2_SO_4_, 8.56 mM NaCl, 2 mM MgSO_4_, 100 μM CaCl_2_, 1 or 10 mM sucrose, pH = 7] containing 50 μg mL^-1^ rifampicin. *Pto* strains were cultured at 28°C for 20 hours with shaking (200 rpm). OD600 was measured using the Implen OD600 right after bacterial cell density adjustment (0-hour post-inoculation, hpi) and 20 hpi.

### Pathogenicity tests of *Pto*

Tomato M82 seeds were germinated in Sunshine Mix # 5 (Sungro), and the resulting seedlings were cultivated under a photoperiod of 12 hours of light and 12 hours of darkness at a temperature of 22 °C and a relative humidity of 60 % for a duration of 21 days. Two days prior to the dip inoculation, *Pto* and its mutant strain *thue* were streaked from glycerol stocks, which had been stored at -80°C, onto KB agar plates supplemented with 50 μg mL^-1^ rifampicin. All strains were incubated at 22 °C for a duration of two days. To prepare the bacterial inoculums at a concentration of 2.5 x 10^7^ colony forming unit (CFU) mL^-1^, the strains were scraped from the KB agar plates and resuspended in 10 mM MgCl_2_. Subsequently, leaves were dipped in bacterial suspension with 0.01 % Silwet-77. Right after inoculation, cotyledons from three individual plants were homogenized in 10 mM MgCl_2_ using a pestle and mortar to create a single biological replicate, with a total of three biological replicates per bacterial genotype. At three days post-inoculation (dpi), true leaves from three individual plants were similarly homogenized in 10 mM MgCl_2_ to produce a single biological replicate, with three biological replicates per bacterial genotype. The homogenized leaf solutions underwent serial dilution (tenfold) in 10 mM MgCl_2_, and 5 μL of each leaf solution was applied to KB agar supplemented with 50 μg mL^-1^ rifampicin. All *Pto* strains were incubated at 22 °C for a duration of 36 hours. To assess the bacterial population present in the dip-inoculated leaves, colonies that formed on the KB agar were counted manually.

## Use of generative AI tools

Claude Opus 5 was used to refine sections of the text for improved readability. Specifically, prompts were used to ask Claude Opus 5 to identify grammatical errors and internal inconsistencies, to shorten or rephrase specified passages, to assess the logical flow of the manuscript. The authors reviewed and verified all output and are fully responsible for the content of the manuscript.

## Acknowledgements

We thank Prof. Jong Hum Kim (Pohang University of Science and Technology, Republic of Korea) for *Pto* and Prof. Choon Tak Kwon (Kyung Hee University, Republic of Korea) for tomato M82. pnCasPA-BEC was a gift from Quanjiang Ji (Addgene plasmid # 113349; http://n2t.net/addgene:113349 ; RRID: Addgene_113349). We also thank the group of Prof. Cho Hongbaek (Sungkyunkwan University, Republic of Korea) for their help with Pto transformation.

## Author contributions

CD: designed and performed the experiments, analyzed the data, and wrote the manuscript; JYK: supervised the project, conceived the study, and wrote the manuscript.

## Data availability

The *Pto* strains that support the findings of this study are available from the corresponding author upon request.

## Supplementary data

The following supplementary data are available at JXB online.

**Fig. S1.**
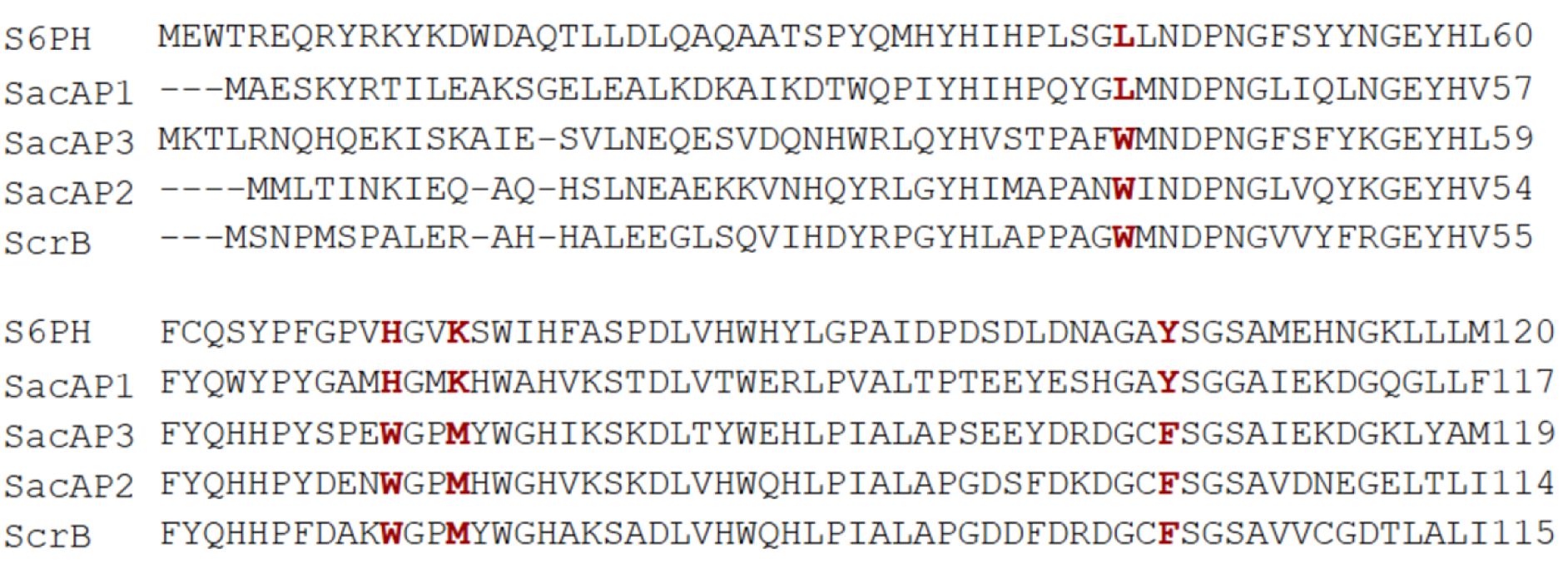
Protein sequence alignment of ScrB showing conserved catalytic residues of sucrose hydrolase. Protein sequence alignment was performed using Clustal Omega (1.2.4) among sucrose-6-phosphate hydrolases from *Lactobacillus gasseri* 224-1 (S6PH) and *Priestia megaterium* (SacAP1), sucrose hydrolases from *Priestia megaterium* (SacAP2 and SacAP3), and ScrB from *Pto*. Key residues that differentiate sucrose hydrolases from sucrose-6-phosphate hydrolases are highlighted in red. Only the partial alignment regions containing these key residues are shown.

**Table S1.**
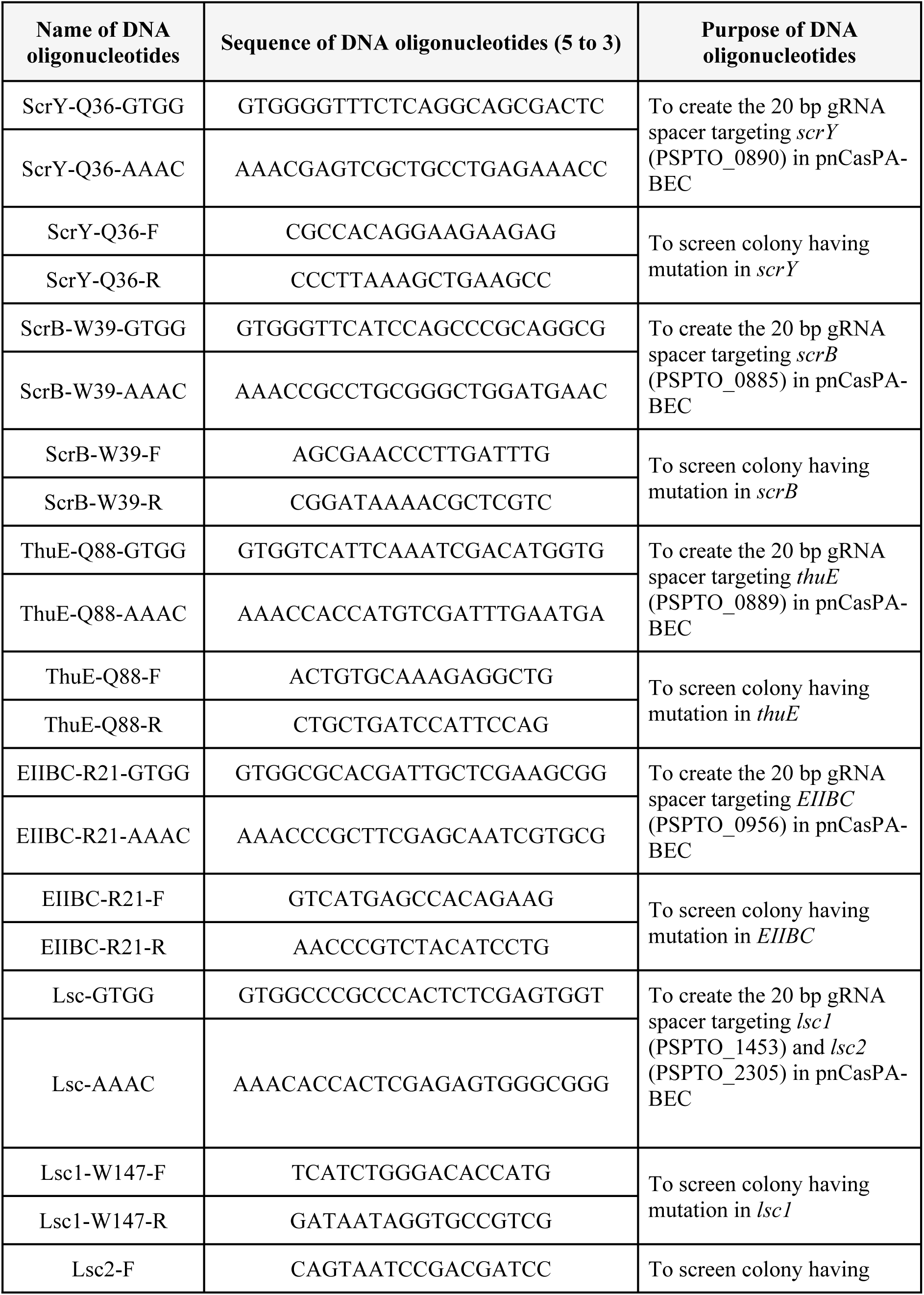
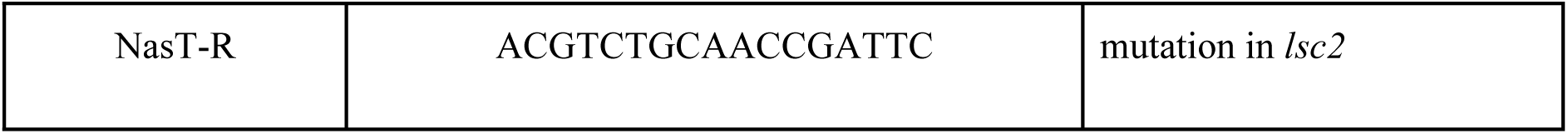
DNA oligonucleotides used in this study

